# The role of calcium, Akt and ERK signaling in cadmium-induced hair cell death

**DOI:** 10.1101/2022.08.15.504004

**Authors:** Jennifer Galdieri, Chloe Adams, María Padilla, Tamara M. Stawicki

## Abstract

Exposure to heavy metals has been shown to cause damage to a variety of different tissues and cell types including hair cells, the sensory cells of our inner ears responsible for hearing and balance. Elevated levels of one such metal, cadmium, have been associated with hearing loss and shown to cause hair cell death in multiple experimental models. While the mechanisms of cadmium-induced cell death have been extensively studied in other cell types they remain relatively unknown in hair cells. We have found that calcium signaling, which is known to play a role in cadmium-induced cell death in other cell types through calmodulin and CaMKII activation as well as IP3 receptor and mitochondrial calcium uniporter mediated calcium flow, does not appear to play a significant role in cadmium-induced hair cell death. While calmodulin inhibition can partially protect hair cells this may be due to impacts on mechanotransduction activity. Removal of extracellular calcium, and inhibiting CaMKII, the IP3 receptor and the mitochondrial calcium uniporter all failed to protect against cadmium-induced hair cell death. We also found cadmium treatment increased pAkt levels in hair cells and pERK levels in supporting cells. This activation may be protective as inhibiting these pathways enhances cadmium-induced hair cell death rather than protecting cells. Thus cadmium-induced hair cell death appears distinct from cadmium-induced cell death in other cell types where calcium, Akt and ERK signaling all promote cell death.

## INTRODUCTION

Hair cells, the sensory cells we use for hearing and balance, are extremely sensitive to insult, and as such disorders of these sensory systems are common. Recently environmental toxins such as heavy metals have gained attention as a potential cause of hearing loss (Choi and Kim, 2014; Schaal et al., 2017). One of these metals is cadmium. Cadmium can be found in some industrial run-off and individuals can be exposed to it in certain occupations, via smoking or via contaminated food (Faroon et al., 2012; IARC, 2012). Multiple studies have found a link between blood or urinary cadmium levels and hearing or balance impairment (Choi et al., 2012; Choi and Park, 2017; Dalton et al., 2020; Huh et al., 2018; Jung, 2019; Kang et al., 2018; Min et al., 2012; Shargorodsky et al., 2011; Wang et al., 2020). Studies have also found hearing loss and/or hair cell death in mammals and fish exposed to cadmium (Faucher et al., 2006; Kim et al., 2008; Liu et al., 2014; Low and Higgs, 2015; Montalbano et al., 2018; Ozcaglar et al., 2001; Schmid et al., 2020; Sonnack et al., 2015). There is evidence that this toxicity may stem from reactive oxygen species generation (ROS). Increased ROS production is seen following treatment with cadmium in the hair cell-like cell line HEI-OC1, and antioxidant treatment protects against cadmium-induced cell death in both HEI-OC1 cells and cochlear explants. Antioxidant treatment also protects against cadmium-induced increases in auditory-brain stem response thresholds in mice (Kim et al., 2008). However, the role of other potential cell-signaling pathways involved in cadmium-induced cell death has not been extensively studied in hair cells to date.

In addition to its association with hearing loss, cadmium has also been linked to a number of other health conditions including cancer, osteoporosis, and dysfunction of the cardiovascular system, kidneys, liver, nervous system and reproductive system. (Bhardwaj et al., 2021; Ferraro et al., 2010; Gallagher et al., 2008; García-Esquinas et al., 2014; Hyder et al., 2013; Mezynska and Brzóska, 2018; Ruczaj and Brzóska, 2022; Tellez-Plaza et al., 2012). While mechanisms of cadmium toxicity in hair cells are still relatively unknown, they have been more extensively studied in other cell types. Multiple cell types show an increase in calcium in response to cadmium (Shih et al., 2005; Smith et al., 1989; Xu et al., 2011; Yamagami et al., 1998; Yang et al., 2007; Yeh et al., 2009). In some cases this increase appears due to the release of calcium from endoplasmic reticulum (ER) stores, as it can be blocked by an inositol 1,4,5-triphosphate (IP3) receptor antagonist but not by the removal of extracellular calcium (Liu et al., 2016; Smith et al., 1989; Yamagami et al., 1998). Cadmium has also been shown to increase IP3 levels in cells (Benters et al., 1997; Hague et al., 2000; Smith et al., 1989; Yamagami et al., 1998). However, in other cases removing extracellular calcium or inhibiting calcium channels can block the cadmium-induced calcium increase suggesting extracellular calcium is contributing to this increase (Liu et al., 2018; Xu et al., 2011; Yeh et al., 2009).

This increase in calcium appears to be important for cadmium-induced cell death as calcium chelation and IP3 receptor inhibition have both been shown to protect against cadmium-induced toxicity (Liu et al., 2016; L. Wang et al., 2009; Wang et al., 2008; Yang et al., 2008, 2007; Yeh et al., 2009). Calcium may ultimately be inducing cell death through disruptions in the mitochondria and the production of ROS as calcium chelation can block cadmium-induced mitochondrial depolarization and increases in ROS production (Xu et al., 2011; Yang et al., 2008, 2007; Yuan et al., 2013). Cadmium is unable to increase ROS production when directly applied to isolated mitochondria further supporting the role of calcium as a potential intermediary signal in this process (Shih et al., 2005). Mitochondrial calcium also appears important in cadmium-induced cell death as inhibition of the mitochondria calcium uniporter blocks cadmium-induced cytochrome c release, mitochondrial depolarization and cell toxicity (Lee et al., 2005; Li et al., 2003; Shih et al., 2005).

Downstream of calcium both calmodulin and calcium/calmodulin-dependent protein kinase II (CaMKII) appear to be playing a role in cadmium-induced cell death. Cadmium is able to directly activate calmodulin as well as increase calcium’s activation of calmodulin, and it is believed this may result in a toxic overactivation of this molecule (Chao et al., 1984; Sutoo et al., 1990). Inhibition of both calmodulin and CaMKII can protect cells against cadmium-induced cell death (Liu et al., 2018; Liu and Templeton, 2007).

Cadmium has also been shown to increase phosphorylated levels of Akt (pAkt), and Extracellular Receptor Kinase (pERK) in vitro in a calcium-dependent manner (Jiang et al., 2015; Xu et al., 2011). Increased pAkt levels have also been seen in the brain in vivo in response to cadmium (Chen et al., 2014). The calcium-dependent activation of these pathways appears to be through the activation of calmodulin and CaMKII as inhabiting either of these molecules can block the cadmium-dependent increase in pAkt and pERK (Chen et al., 2011; Fujiki et al., 2013; Liu et al., 2018; Xiao et al., 2009; Xu et al., 2011). Akt and ERK activation appears to be playing a role in the cell death process as inhibitors of both signaling pathways have been shown to protect against cadmium-induced cell toxicity (Chen et al., 2008; Iryo et al., 2000; Jiang et al., 2014, 2015; Y. K. Kim et al., 2005; Wang et al., 2014; Zhang et al., 2017; Zhao et al., 2015).

Calcium, Akt and ERK signaling have also all been previously implicated in hair cell damage from either aminoglycoside antibiotic toxicity or noise. Inhibiting calcium movement from the ER or mitochondria has been shown to protect against hair cell damage (Esterberg et al., 2014; Wang et al., 2019). There is evidence that a transfer of calcium from the ER to the mitochondria in response to aminoglycoside treatment is triggering ROS generation leading to hair cell death (Esterberg et al., 2016). The Akt and ERK signaling pathways on the other hand are believed to play a protective role in hair cell toxicity as inhibition of these pathways is most commonly shown to increase hair cell damage (Battaglia et al., 2003; Brand et al., 2015; Chung et al., 2006; Haake et al., 2009; Hayashi et al., 2013; Kurioka et al., 2015). Therefore, we felt these three signaling molecules were strong candidates to investigate as potential signaling pathways involved in cadmium-induced hair cell death. Activation of ERK has also previously been seen in the cochlea in response to cadmium and inhibition of ERK can protect HE-OC1 cells, a hair cell like cell line, from cadmium-induced cell death (Kim et al., 2008).

We investigated the role of these signaling molecules in hair cell death using the zebrafish lateral line. The lateral line is a sensory system in fish used to detect water motion that consists of clusters of hair cells and support cells, collectively known as neuromasts, on the surface of the animal. Support cells are glia-like cells surrounding hair cells that have been shown to play key roles in the development, maintenance and regeneration of hair cells (reviewed in Monzack and Cunningham, 2013). Hair cells of the zebrafish lateral line have been shown to be functionally, molecularly and morphologically similar to mammalian hair cells (reviewed in Nicolson, 2005; Whitfield, 2002).

We have previously shown that cadmium causes reliable hair cell death in the zebrafish lateral line within three hours in a dose-dependent manner, and therefore used that treatment paradigm in this study (Schmid et al., 2020). We first found that removal of extracellular calcium following cadmium treatment caused an increase in hair cell death both in control conditions and following cadmium treatment. We also found that co-treatment of a calmodulin inhibitor with cadmium could significantly protect against cadmium-induced hair cell death. However, when the calmodulin inhibitor was applied after fish were washed out of cadmium, a condition we refer to as posttreatment, protection was no longer seen. Additionally, other calcium signaling pathway inhibitors including a CaMKII inhibitor, inositol IP3 receptor inhibitor, and mitochondria calcium uniporter inhibitor failed to protect against cadmium-induced hair cell death. We also saw an increase in levels of pAkt in hair cells and pERK in supporting cells in response to cadmium, however, inhibition of these pathways enhanced hair cell death rather than protecting against it. Therefore, we conclude that the signaling pathways responsible for cadmium-induced hair cell death appear distinct from those commonly seen in cadmium-induced toxicity in other cell types and partially distinct from those seen in other forms of hair cell toxicity.

## MATERIALS AND METHODS

### Animals

All experiments were carried out in 5 days post-fertilization (dpf) *AB strain wild-type zebrafish larvae. Fish larvae were housed in an incubator maintained at 28.5°C with a 14/10 hour light/dark cycle and raised in embryo media (EM) consisting of 1 mM MgSO_4_, 150 μM KH_2_PO_4_, 42 μM Na_2_HPO_4_, 1 mM CaCl_2_, 500 μM KCl, 15 mM NaCl, and 714 μM NaHCO_3_. The Lafayette College Institution Animal Care and Use Committee approved all experiments.

### Drug Treatments

The following drugs were used: cadmium chloride hemipentahydrate (Fisher Scientific), water-soluble KN-93 (Abcam), MK-2206 (Millipore Sigma), PD184352 (Millipore Sigma), RU360 (Millipore Sigma), W-13 (Enzo) and Xestospongin (Abcam). Stock solutions for cadmium chloride, KN-93, and Ru360 were made in water. Stock solutions of all other drugs were made in dimethyl sulfoxide (DMSO, Millipore Sigma). All drugs were then diluted to their treatment concentrations in EM. For drugs diluted in DMSO the control groups contained DMSO at the same volume as the drug. For the water diluted drugs EM was used for the control group.

For the drug treatments, fish were initially put into netwell inserts (Corning) in 6 well plates containing EM. The netwell inserts with the larvae were then moved to new plates containing the solutions the fish were to be treated in. At the end of treatment, fish were washed three times in EM, again by moving the netwell inserts to new plates, between treatments or before fixation. For experiments where fish were moved to EM with reduced calcium levels after cadmium treatment, they were washed three times in zero calcium EM instead of standard EM before being moved to the reduced or zero calcium EM and then returned to standard EM immediately prior to fixation.

### Immunostaining

Fish were anesthetized with MS-222 and then fixed for 2 hours at room temperature in 4% paraformaldehyde. Antibody labeling was carried out as previously described (Stawicki et al., 2014). The following primary antibodies were used: rabbit anti-parvalbumin (ThermoFisher, PA1-933) diluted at 1:1,000, mouse anti-otoferlin (Developmental Studies Hybridoma Bank, HCS-1) diluted at 1:100, rabbit anti-phospho-Akt (Ser473) (Cell Signaling, 9271) diluted at 1:100, and rabbit anti-phospho-p44/42 MAPK (ERK1/2) (Thr202/Tyr204) (Cell Signaling, 4370) diluted at 1:200.

### Hair Cell Counts

For hair cell counts fish were either stained with just parvalbumin or costained with otoferlin and parvalbumin. In the majority of experiments the parvalbumin stain was used for counts, however, in fish that were costained with otoferlin the otoferlin stain was used if the parvalbumin stain was too dim to reliably count hair cells. Within a single experiment the same antibody was always used for all counts. To perform hair cell counts fish were observed on an Accu-Scope EXC-350 microscope using the 40X objective. The number of hair cells in the following neuromasts OP1, M2, IO4, O2, MI2, and MI1 (Raible and Kruse, 2000) were counted for each larvae. The total number of hair cells counted for that fish was then divided by 6 to get an average number of hair cells per neuromast. To normalize data for the zero calcium and ERK inhibitor experiment, the average hair cells/neuromast number for each treatment group was divided by the average hair cells/neuromast number for their respective control groups with no cadmium and then multiplied by 100 to be converted to a percentage.

### Confocal Imaging and Analysis of pAkt and pERK levels

To calculate pAkt and pERK levels fish stained with either the pAkt or pERK antibodies and the otoferlin antibody were imaged on a Zeiss LSM800 confocal microscope using the Zen Blue software. For pAkt, the approximate center of the neuromast was selected and 10 planes separated by 1 μm were imaged for each neuromast. For pERK, there was significant neuronal labeling in the control condition that we did not want to include in our analysis. Therefore, the top and bottom of the neuromast were selected and planes separated by 1μm were imaged throughout the neuromast regardless of number with images ranging from 11-29 planes. Then 10 planes with minimal neuronal labeling were selected for subsequent analysis. One neuromast of the anterior lateral line was imaged per fish with the exact neuromast to be imaged selected based on which neuromasts were best positioned to image. Image analysis was conducted in Fiji. Maximum projection images were made for each 10-plane stack. The neuromast was outlined using the otoferlin label and the average fluorescent intensity of the pAkt or pERK signal was measured.

### Statistics and Graphical Representation of Data

All experiments were carried out with n=10 animals per group. Statistics were calculated in GraphPad Prism 6 which was also used to generate graphs. For experiments where two or more different groups were tested against a range of calcium or cadmium doses, a two-way ANOVA was used with the either a Tukey’s or Šídák’s multiple comparisons test depending on which was recommended by Prism. The multiple comparisons test used for each experiment is mentioned in the figure legend. Graphs of this data were represented by the average of each data set with error bars showing the standard deviation. For all other experiments, a one-way ANOVA was used again with a Tukey’s multiple comparison test as all groups were compared. Graphs of this data show the individual data points as well as the average and standard deviations. For experiments where the ANOVA showed a significant difference between relevant groups the significance found by the multiple comparisons tests are shown on the graphs as asterisks.

## RESULTS

### Alterations in extracellular calcium levels following cadmium treatment do not cause dramatic changes in cadmium-induced hair cell death

Previous research has shown that calcium plays an important role in cadmium-induced cell death in many cell types. In some cell types, it appears that extracellular calcium is important as chelating it or treating cells with calcium channel blockers can protect against cadmium-induced toxicity (Liu et al., 2018; Yang et al., 2008; Yeh et al., 2009). Unfortunately, determining the role of extracellular calcium in cadmium-induced hair cell toxicity is complicated by the fact that we have previously shown that cadmium-induced hair cell toxicity is mechanotransduction-activity dependent (Schmid et al., 2020). Extracellular calcium levels are known to impact hair cell mechanotransduction activity (reviewed in Fettiplace, 2017; Ó Maoiléidigh and Ricci, 2019) with calcium chelation completely blocking it (Assad et al., 1991). It has previously been shown that the entry and toxicity of other mechanotransduction-dependent ototoxins can be altered by alterations in extracellular calcium levels for this reason (Coffin et al., 2009). Therefore, to investigate whether calcium signaling was impacting cadmium-induced hair cell death we first wanted to determine if there was continued hair cell death following cadmium removal. If there was, we could potentially separate cadmium entry, which is the phase of cadmium-induced hair cell death most likely to be mechanotransduction dependent, from the subsequent cell death process. To do this we treated fish with 30 μM of cadmium chloride for either 1 or 2 hours, fish were then washed out of cadmium, and either fixed immediately or fixed three hours after the start of the experiment. We found that following both the 1 and 2 hour cadmium chloride treatment continued hair cell death was seen after cadmium removal (Figure 1).

**Figure 1:**
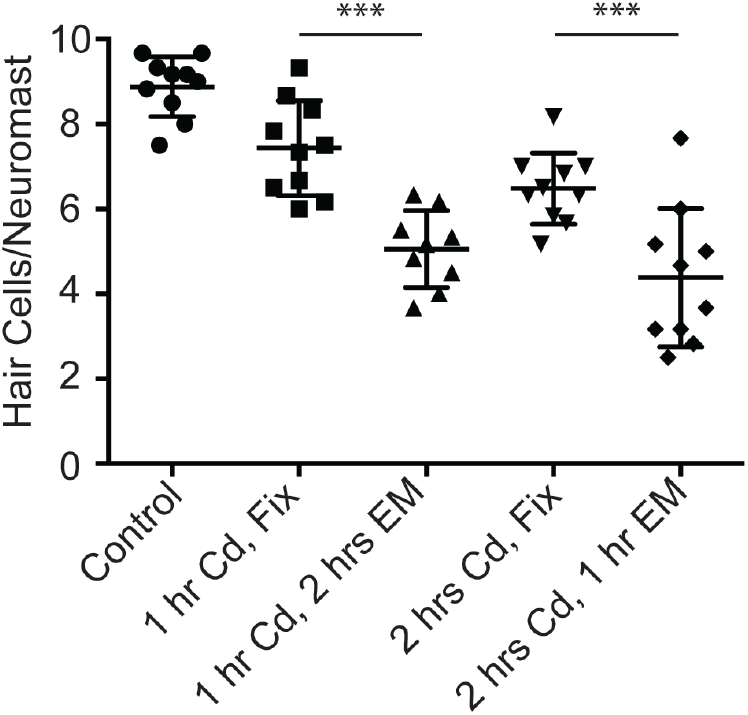
Cadmium causes continued cell death after removal. Hair cells remaining after various treatment paradigms with 30μM of cadmium. Fish were either treated with cadmium chloride for 1 hour and immediately fixed (squares), treated with cadmium for 1 hour then left in EM for two hours before fixation (triangles), treated with cadmium for 2 hours and then immediately fixed (upside down triangles), or treated with cadmium for 2 hours and then left in EM for 1 hour before fixation (diamonds). Control fish were kept in EM for three hours (circles). There was a significant amount of cell death after cadmium removal for both the 1 hour and 2 hours cadmium treatment paradigms (*** = p<0.001 by ANOVA and Tukey post-hoc test).

As we saw significant continued cadmium-induced hair cell death after cadmium removal, we next wanted to test if this continued cell death could be altered by changing extracellular calcium levels. To do this we used the treatment paradigm where fish were treated with cadmium chloride for one hour, washed out of cadmium, and then fixed two hours later. We first tested a range of calcium concentrations by treating fish with either EM or 30 μM cadmium chloride for 1 hour and then washing them out of that solution and moving them into EM with calcium concentrations ranging from 0-2 mM for 2 hours prior to fixation. We found the only calcium concentration where a significant difference was seen in the amount of cadmium-induced hair cell death as compared to the standard EM calcium concentration of 1mM was 0mM (Figure 2A). However, rather than protecting hair cells the removal of extracellular calcium appeared to increase cadmium-induced hair cell death. We then treated fish with a range of cadmium doses from 0-120 μM for 1 hour before washing them out of cadmium and moving them to either standard EM with 1 mM calcium or EM with 0 mM calcium for 2 hours prior to fixation. Again, a significant increase in the amount of cadmium-induced hair cell death was seen in the fish that were moved to 0 mM calcium following cadmium treatment (Figure 2B). However, there was also a significant reduction in the number of control hair cells in this condition similar to what has previously been seen when zebrafish are put in calcium-free EM (Coffin et al., 2009). Therefore, to determine whether the incubation in 0 mM calcium EM was sensitizing fish to cadmium-induced hair cell death or simply killing hair cells on its own we normalized these data for both the 1 mM and 0 mM calcium EM groups to the 0 cadmium control for each group. We found in this case while there is still a trend toward a decrease in hair cell number at all cadmium concentrations tested, the only concentration that showed a significant decrease was 120 μM cadmium (Figure 2C). Taken together this data suggests alterations in calcium concentration following cadmium removal do not have dramatic impacts on cadmium-induced hair cell death. However, due to the complications with calcium’s effect on mechanotransduction activity, we cannot rule out an impact from extracellular calcium at earlier time points in the cell death process.

**Figure 2:**
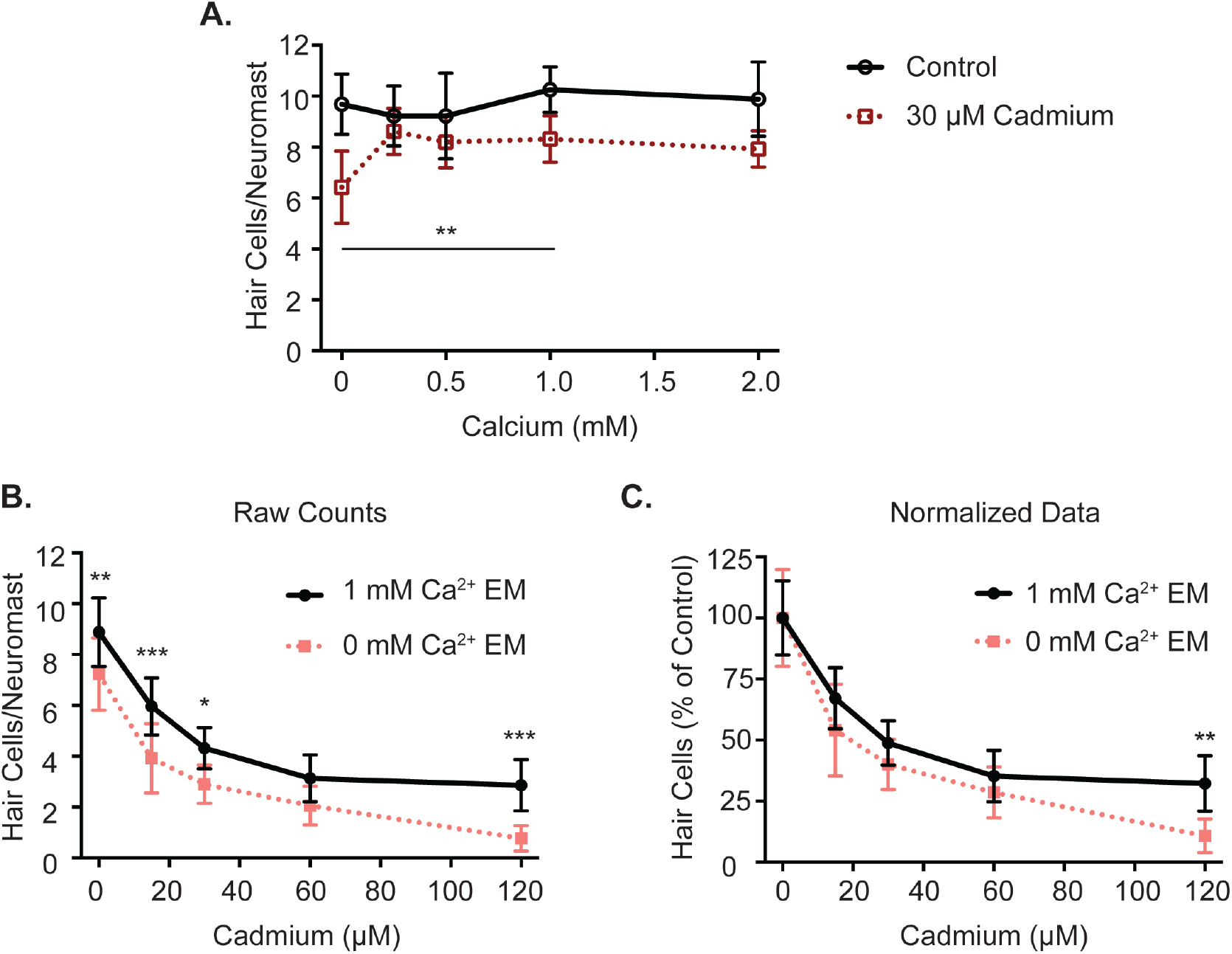
Removal of extracellular calcium following cadmium treatment kills hair cells on its own but does not dramatically impact cadmium-induced hair cell death. (A) Hair cells remaining after a 1 hour treatment with either EM, control (solid line), or 30 μM cadmium chloride (dotted line) followed by a 2 hour incubation in EM containing varying concentrations of calcium. The only calcium concentration where there was a significant difference in the number of hair cells following cadmium treatment compared to the standard EM 1 mM calcium group was the 0 mM calcium group (** = p<0.01 by Two-Way ANOVA and Tukey’s post-hoc test). (B) Hair cells remaining after a 1 hour treatment with varying doses of cadmium chloride followed by a 2 hour incubation in either EM containing 1mM calcium (solid line) or EM containing 0 mM calcium (dotted line). A significant decrease in hair cells in the 0 mM calcium group was seen for all concentrations of cadmium tested except 60 μM (* = p<0.05, ** = p<0.01, *** = p<0.001, by Two-Way ANOVA and Šídák’s post-hoc test). (C) The same data as in B, but normalized to the zero cadmium control for each group. Following normalization, a significant decrease in hair cells in the 0 mM calcium group was only seen at 120 μM cadmium. (** = p<0.01, by Two-Way ANOVA and Šídák’s post-hoc test). N = 10 fish for each group. Error bars = Standard deviation.

### Calmodulin and CaMKII inhibitors do not consistently protect against cadmium-induced hair cell death

Calmodulin is a major intracellular receptor for calcium and calmodulin inhibition has been shown to protect against cadmium-induced cell death (Niewenhuis and Prozialeck, 1987; Xu et al., 2011; Yang and Yang, 1997). Calmodulin inhibition has also previously been shown to protect against aminoglycoside-induced hair cell death (Esterberg et al., 2013). To see if calmodulin inhibition could protect against cadmium-induced hair cell death we first cotreated fish with 30 μM of the calmodulin inhibitor W-13 hydrochloride and varying doses of calmodulin chloride ranging from 0-120 μM for three hours. We found that calmodulin inhibition significantly reduced cadmium-induced hair cell death at lower doses of cadmium chloride, 15 and 30 μM, but not at higher doses, 60 and 120 μM (Figure 3A).

**Figure 3:**
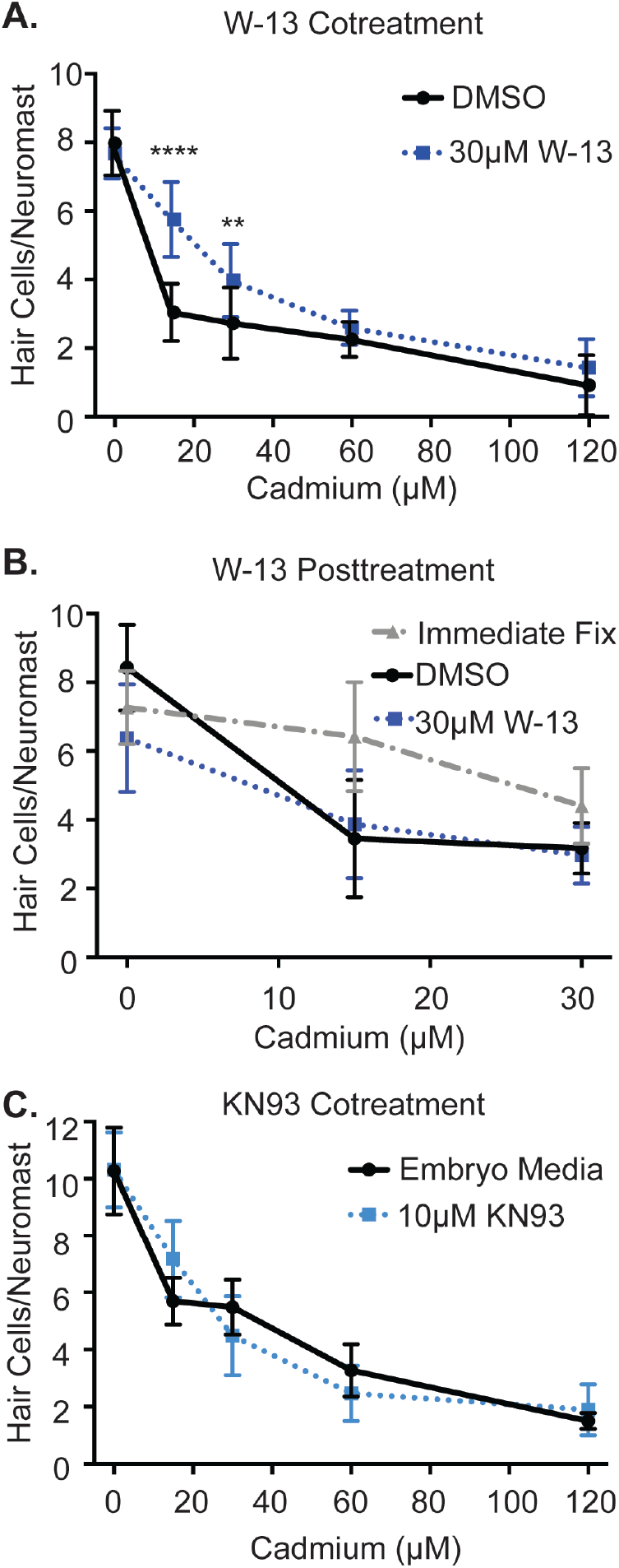
Calmodulin inhibitor cotreatment protects against cadmium-induced hair cell death whereas calmodulin inhibitor posttreatment and CaMKII inhibitor cotreatment fail to protect. (A) Hair cells remaining after a 3 hour cotreatment of either DMSO, a vehicle control, (solid line), or 30 μM of the calmodulin inhibitor W-13 (dotted line) and varying doses of cadmium chloride. W-13 cotreatment caused a significant reduction in hair cell death seen at lower cadmium doses (** = p<0.01, **** = p<0.0001 by Two-Way ANOVA and Šídák’s post-hoc test). (B) Hair cells remaining after a 1 hour treatment with varying doses of cadmium chloride and either immediate fixation (dashed line), or a two hour post-treatment with DMSO (solid line) or W-13 (dotted line). No significant protection was seen comparing the DMSO and W-13 groups by Two-Way ANOVA and Tukey’s post-hoc test. (C) Hair cells remaining after a 3 hour cotreatment of either plain embryo media or 10 μM of the CaMKII inhibitor KN-93 and varying doses of cadmium chloride. No significant difference was seen between the two groups by Two-Way ANOVA. n = 10 fish for each group. Error bars = Standard deviation.

However, calmodulin inhibition has also previously been shown to inhibit rapid apical endocytosis in hair cells as measured by rapid FM1-43 uptake, a process dependent on mechanotransduction activity (Seiler and Nicolson, 1999). As mentioned above inhibiting mechanotransduction can protect against cadmium-induced hair cell death (Schmid et al., 2020), therefore, we tested whether inhibiting calmodulin only after cadmium removal would still significantly protect against cadmium-induced hair cell death. To do this fish were treated with varying doses of cadmium chloride from 0-30 μM for one hour, washed out of cadmium, and then immediately fixed or placed into DMSO or 30 μM W-13 for two hours before fixation. Using this treatment paradigm we no longer saw protection from calmodulin inhibition (Figure 3B). This suggests the protection seen with cotreatment with the calmodulin inhibitor may be due to impacts on mechanotransduction-dependent processes, though we can also not rule out a distinct early role for calmodulin in the cell death process.

Calmodulin is known to activate CaMKII and inhibiting CaMKII has also been shown to protect against cadmium-induced cell death in multiple cell types (Chen et al., 2011; Liu et al., 2018; Liu and Templeton, 2007). Therefore, as our calmodulin results did now show a clear protective impact we, subsequently tested if CaMKII inhibition would impact cadmium-induced hair cell death. To do this we cotreated fish with 10 μM of the CaMKII inhibitor KN-93, a dose previously shown to be effective in zebrafish (Rothschild et al., 2013), and varying doses of cadmium chloride for three hours. We found CaMKII inhibition did not significantly impact cadmium-induced hair cell death (Figure 3C). This further supports the idea that calmodulin and CaMKII are not playing a role in cadmium-induced hair cell death outside of calmodulin’s impact on mechanotransduction activity.

### Inhibition of the IP3 Receptor and Mitochondrial Uniporter fail to protect against cadmium-induced hair cell death

We next wanted to test whether calcium flow from the ER or mitochondria would impact cadmium-induced hair cell death as blocking the IP3 receptor and the mitochondria calcium uniporter has been shown to protect other cell types against cadmium-induced toxicity (Lee et al., 2005; Liu et al., 2016, 2018; Shih et al., 2005; S. H. Wang et al., 2009; Wang et al., 2008, 2022). These inhibitors have also been shown to protect hair cells from other forms of toxicity. (Esterberg et al., 2014; Wang et al., 2019). To test this in hair cells we cotreated fish for three hours with either 1 μM of the IP3 receptor inhibitor Xestospongin C or 500 nM of the mitochondrial uniporter inhibitor RU360, doses that have previously been shown to protect against aminoglycoside-induced hair cell death in the zebrafish lateral line (Esterberg et al., 2014), and varying doses of cadmium chloride for three hours. In both cases, the inhibitor failed to significantly protect against cadmium-induced hair cell death with RU360 causing a slight enhancement of cadmium-induced hair cell death at the lowest cadmium dose (Figure 4). Thus, our results indicate calcium flow from the ER and mitochondria is not playing a significant role in cadmium-induced hair cell death.

**Figure 4:**
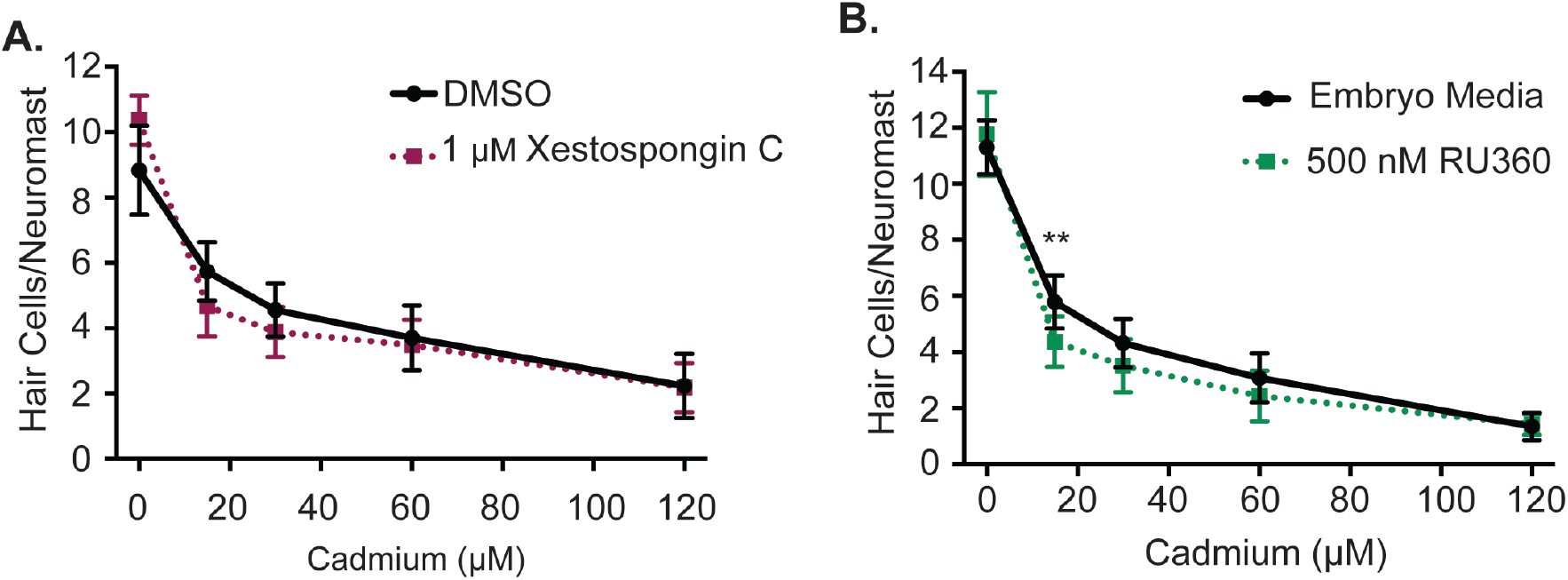
Inhibiting ER and mitochondria calcium transfer does not protect against cadmium-induced hair cell death. Hair cells remaining after a three hour cotreatment of either EM or DMSO (solid lines), or (A) 1 μM of the IP3 receptor inhibitor Xestospongin C, or (B) 500 nM of the mitochondrial uniporter inhibitor RU360 (dotted lines) and varying doses of cadmium chloride. Neither of the inhibitors significantly protected against cadmium-induced hair cell death with RU360 showing a significant increase in cadmium-induced hair cell death at the lowest dose (** = p<0.01= by Two-Way ANOVA and Šídák’s post-hoc test). N = 10 fish/group. Error bars = standard deviation.

### Cadmium increases Akt and ERK signaling in neuromasts

In many cell types cadmium increases the level of phosphorylated Akt and ERK (Chen et al., 2008, 2011; Fujiki et al., 2013; Iryo et al., 2000; J. Kim et al., 2005; Kim et al., 2008; Liu and Templeton, 2007; Misra et al., 2003; Xiao et al., 2009; Xu et al., 2011). Activation of these pathways has also been seen in hair cells or supporting cells in response to other forms of hair cell damage (Chung et al., 2006; Herranen et al., 2018; Ingersoll et al., 2020; Lahne and Gale, 2008; Maeda et al., 2013). Based on this evidence we tested whether the Akt or ERK pathways would be activated in either hair cells or supporting cells of the zebrafish lateral line in response to cadmium. To do this we treated fish with 30 μM of cadmium chloride for one hour and then fixed and stained with an antibody for either pAkt or pERK. We found that treatment with cadmium significantly increased the levels of pAkt seen in hair cells (Figure 5A & B). This increase was lost when fish were cotreated with cadmium and 5 μM of the Akt inhibitor MK-2206. We also saw a significant increase in pERK levels following cadmium treatment that was lost when fish were cotreated with cadmium and 3 μM of the MEK inhibitor PD184352 which suppresses ERK phosphorylation (Figure 5C & D). However, unlike pAkt labeling which appeared localized to hair cells, pERK labeling appears localized to supporting cells (Figure 5B & D). This is in agreement with past research in mammals showing increased pERK in support cells but not hair cells in response to other forms of hair cell damage (Herranen et al., 2018; Ingersoll et al., 2020; Lahne and Gale, 2008; Maeda et al., 2013).

**Figure 5:**
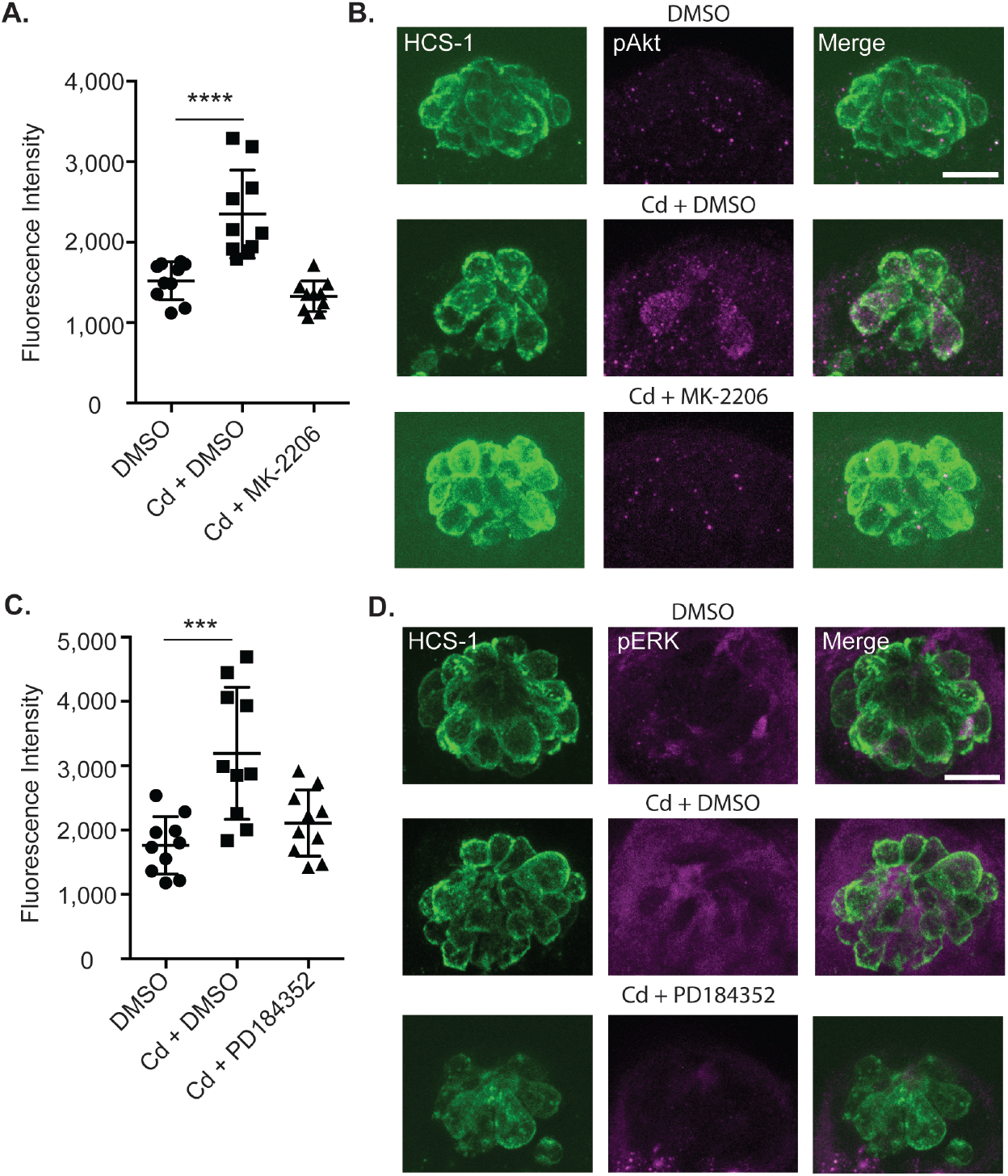
Cadmium treatment increases pAkt levels in hair cells and pERK levels in supporting cells. (A) Quantification of the fluorescence intensity of pAkt in fish treated for 1 hour with either DMSO alone, 30μM cadmium chloride (Cd) + DMSO, or 30μM Cd and 5 μM of MK-2206. (**** = p<0.0001 by ANOVA and Tukey post-hoc test). (B) Images of a neuromast from the DMSO alone group (top), the Cd + DMSO group (middle), and Cd + MK-2206 group (bottom). Hair cells are labeled with the otoferlin (HCS-1) antibody in green (left) and pAkt in magenta (center). (C) Quantification of the fluorescence intensity of pERK in fish treated for 1 hour with either DMSO alone, 30μM Cd + DMSO, or 30μM Cd and 3 μM of PD184352. (*** = p<0.001 by ANOVA and Tukey post-hoc test). (D) Images of a neuromast from the DMSO alone group (top) and the Cd + DMSO group (middle), and Cd + MK-2206 group (bottom). Hair cells are labeled with the HCS-1 antibody in green (left) and pERK in magenta (center). Scale bar = 10 μm.

### Inhibiting Akt and ERK signaling pathways increases cadmium-induced hair cell death

Having seen an increase in pAkt and pERK levels following cadmium treatment, we wanted to determine whether the Akt or ERK signaling pathways impacted cadmium-induced hair cell death. Akt and/or ERK inhibitors have been shown to protect against cadmium-induced toxicity in multiple cell types (Chen et al., 2008; Iryo et al., 2000; Jiang et al., 2015; J. Kim et al., 2005; Kim et al., 2008; Wang et al., 2014; Zhang et al., 2017), suggesting these pathways normally function to promote cadmium-induced cell death. In hair cells blocking Akt or ERK signaling commonly increases hair cell death, suggesting these pathways may be activated to protect hair cells (Battaglia et al., 2003; Brand et al., 2015; Chung et al., 2006; Haake et al., 2009; Hayashi et al., 2013; Kurioka et al., 2015).

To determine if the Akt or ERK pathways were impacting cadmium-induced hair cell death, we cotreated fish with either 5 μM of the Akt inhibitor MK-2206, or 1.5 μM of the MEK inhibitor PD184352, which inhibits ERK activation, and varying doses of cadmium chloride for three hours. A smaller dose of PD184352 was used in this experiment than the pERK levels experiment due to increased hair cell toxicity from PD184352 alone with the longer treatment time. We found both drugs caused a decrease in hair cell number following cadmium treatment particularly, at the 30 and 60 μM cadmium chloride doses (Figure 6A & B). However, as noted above PD184352 caused a loss of hair cells independent of cadmium treatment, a similar loss of hair cells from simple ERK inhibition has also previously been seen in cochlear explants (Battaglia et al., 2003). Therefore, to determine if PD184352 was enhancing cadmium-induced hair cell death or simply killing hair cells on its own, we normalized the data for both the DMSO and PD184352 groups to the zero cadmium control (Figure 6C). In doing so we still saw a significant increase in cadmium-induced hair cell death at the 30 and 60 μM cadmium chloride doses. Thus, we conclude that unlike in other cell types Akt and ERK signaling do not appear to be facilitating cadmium-induced hair cell death and may be playing a protective role similar to what is seen in other forms of hair cell toxicity.

**Figure 6:**
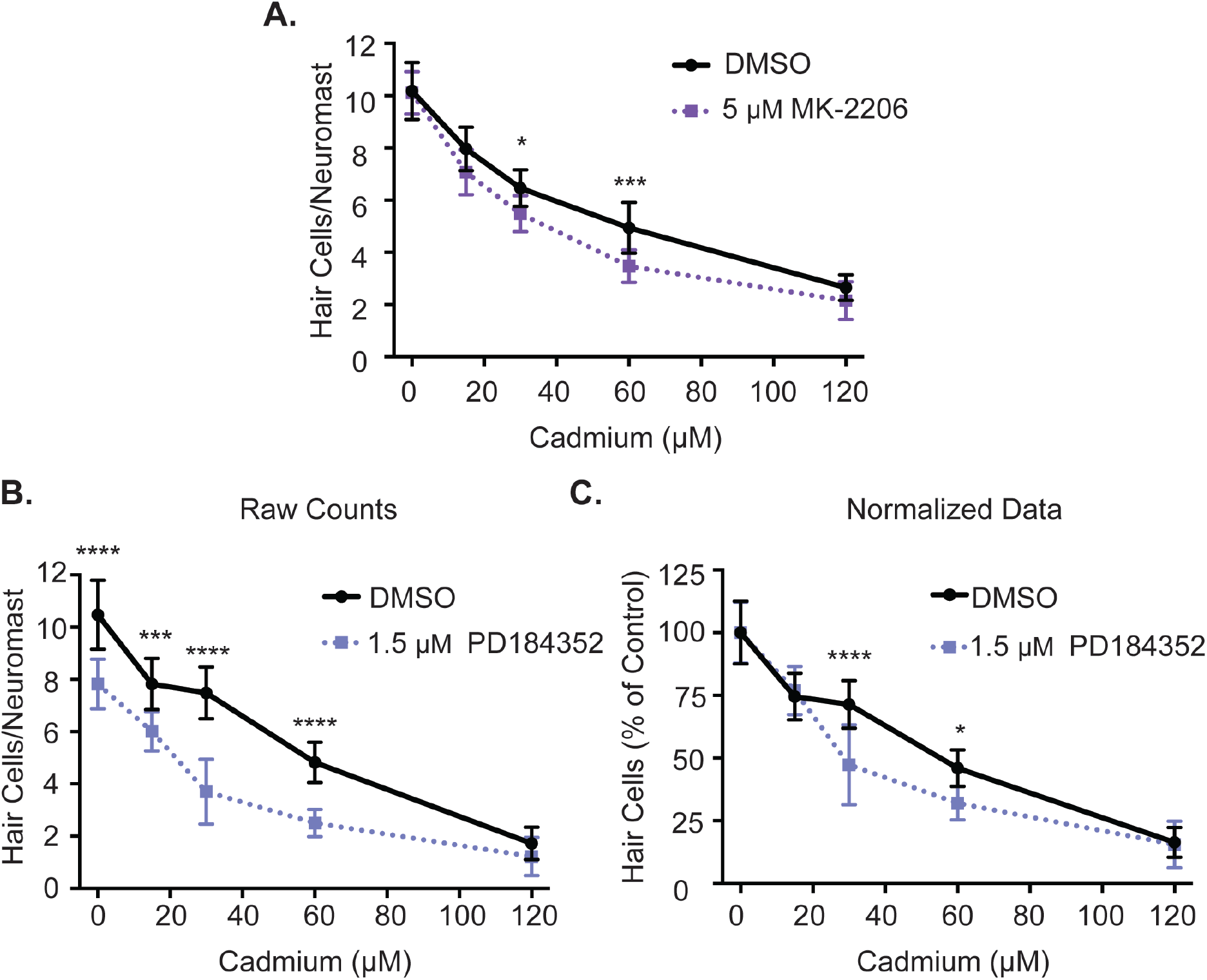
Inhibiting Akt and ERK signaling increases cadmium-induced cell death. (A) Hair cells remaining after a 3 hour cotreatment of either DMSO (solid line) or 5 μM of the Akt inhibitor MK-2206 (dotted line) and varying doses of cadmium chloride. MK-2206 cotreatment caused a significant increase in cadmium-induced hair cell death. (B) Hair cells remaining after a 3 hour cotreatment of either DMSO, a vehicle control, (solid line), or 1.5 μM of the ERK pathway inhibitor PD184352 (dotted line) and varying doses of cadmium chloride. PD184352 caused a significant decrease in the number of hair cells seen. (C) The same data as in B, but normalized to the zero cadmium control for each group. PD184352 caused a significant increase in the amount of cadmium-induced hair cell death seen. (* = p<0.05, *** = p<0.001, **** = p<0.0001, by Two-Way ANOVA and Šídák’s post-hoc test). N = 10 fish for each group. Error bars = Standard deviation.

## DISCUSSION

Overall we found that cadmium-induced hair cell death showed both some distinctions and similarities with cadmium-induced cell death in other cell types and other forms of hair cell toxicity. One of the distinctions was that we failed to find a significant role for calcium signaling. This is in contrast to both cadmium-induced cell death in other cell types and other forms of hair cell death where calcium appears to be a key mediator in the cell death process (Esterberg et al., 2014, 2013; Liu et al., 2016; L. Wang et al., 2009; Wang et al., 2008, 2019; Xu et al., 2011; Yang et al., 2008, 2007; Yeh et al., 2009). We did however, see activation of both the Akt and ERK signaling pathways. These two pathways are known to be activated both in response to cadmium in other cell types (Chen et al., 2008, 2011; Fujiki et al., 2013; Iryo et al., 2000; J. Kim et al., 2005; Kim et al., 2008; Liu and Templeton, 2007; Misra et al., 2003; Xiao et al., 2009; Xu et al., 2011) and in other forms of hair cell toxicity (Chung et al., 2006; Herranen et al., 2018; Ingersoll et al., 2020; Lahne and Gale, 2008; Maeda et al., 2013). Though unlike in other cell types, inhibiting these signaling pathways enhanced cadmium-induced hair cell death rather than protecting against it (Chen et al., 2008; Iryo et al., 2000; Jiang et al., 2014, 2015; Y. K. Kim et al., 2005; Wang et al., 2014; Zhang et al., 2017; Zhao et al., 2015) similar to what has been shown in other forms of hair cell toxicity (Battaglia et al., 2003; Brand et al., 2015; Chung et al., 2006; Haake et al., 2009; Hayashi et al., 2013; Kurioka et al., 2015).

While most of our manipulations of calcium signaling failed to protect against cadmium-induced hair cell death, one manipulation that did cause significant protection was the cotreatment of cadmium and a calmodulin inhibitor. It has been shown that cadmium can activate calmodulin in lieu of calcium and it has been hypothesized this may be one potential mechanism of cadmium-induced toxicity (Chao et al., 1984; Sutoo et al., 1990). Treatment with inhibitors blocking calmodulin-dependent signaling pathways has been shown to protect against cadmium in multiple cell types and tissues including the kidneys, osteoblasts, and testes (Ha et al., 2016; Niewenhuis and Prozialeck, 1987; Yang and Yang, 1997). However, these experiments are complicated in hair cells by the fact that calmodulin inhibition also inhibits rapid apical endocytosis in hair cells, a process dependent on mechanotransduction activity (Seiler and Nicolson, 1999). As inhibiting mechanotransduction activity is known to inhibit cadmium-induced hair cell death (Schmid et al., 2020) we cannot definitively say that the protection we see from calmodulin inhibition is due to a role for calmodulin in the cell death process. The fact that protection is no longer seen when the calmodulin inhibitor is added after cadmium treatment, rather than as a cotreatment, and the fact that CaMKII inhibition was unable to protect hair cells from cadmium-induced hair cell death further supports the idea that calmodulin inhibition is mainly impacting hair cell mechanotransduction activity and cadmium entry rather than the cell death process.

We also failed to see protection against cadmium-induced hair cell death using other calcium signaling manipulations including removing extracellular calcium and inhibiting the IP3 receptor and mitochondrial calcium uniporter. These manipulations have been shown to protect against cadmium-induced cell death in other cell types (Lee et al., 2005; Liu et al., 2016, 2018; Shih et al., 2005; Wang et al., 2008; Xu et al., 2011). However, these previous studies were carried out in mitotic cultured cells, including mesangial cells, PC-12 cells, proximal tubule cells, osteoblasts, and osteosarcoma cells, whereas our study was carried out *in vivo* in postmitotic cells, so this may be the reason for the discrepancy. Indeed another study looking at neurons, another postmitotic cell, failed to see protection from cadmium-induced cell death when inhibiting the mitochondrial calcium uniporter (Tang et al., 2021). Calcium signaling in general and specifically calmodulin, calcium release via the IP3 receptor, and activation of the mitochondrial uniporter have all been shown to be important for the progression of mitosis (Lagos-Cabré et al., 2020; Nugues et al., 2022; Rasmussen and Means, 1989; Takuwa et al., 1995; Zhao et al., 2019). Therefore, mitotic cells may be more sensitive to cadmium-induced alterations in these calcium signaling pathways. Also, even in mitotic cells, there have been discrepancies over the role of calcium signaling in cadmium-induced cell death. For example, when looking at CaMKII inhibition a study in an osteoblast cell line showed no effect (Ha et al., 2016), and a study in mesangial cells showed an increase in cell death (Xiao et al., 2009).

The fact that blocking the IP3 receptor and mitochondrial calcium uniporter failed to protect against cadmium-induced hair cell death also contrasted with what has previously been shown in other forms of hair cell death (Esterberg et al., 2014; Wang et al., 2019). However, while there are clearly some overlapping mechanisms between different forms of hair cell death, drug screens frequently find drugs that can protect against only a subset of ototoxic insults (Coffin et al., 2013; Owens et al., 2009; Vlasits et al., 2012) showing there are distinct mechanisms by which ototoxic compounds induce cell death in hair cells.

While we did not find a significant impact of calcium signaling in cadmium-induced hair cell death we did see an increase in Akt and ERK signaling similar to what has been seen in other cell types. This was somewhat unexpected as in other cell types this increase is calcium signaling dependent (Chen et al., 2011; Fujiki et al., 2013; Jiang et al., 2015; Liu et al., 2018; Xiao et al., 2009; Xu et al., 2011). However, it has previously been shown that ERK signaling can be activated in supporting cells even following calcium chelation (Lahne and Gale, 2008). One potential alternative mechanism for the activation of these pathways includes Ras and Raf signaling which has previously been implicated in activating ERK signaling following hair cell damage (Battaglia et al., 2003; Lahne and Gale, 2008). Another alternative activation route is growth factor signaling. Increases in growth factor receptors have been seen after hair cell damage (Lee and Cotanche, 1996; Zine and de Ribaupierre, 1999), and inhibiting the IGF1R receptor has been shown to also increase susceptibility to aminoglycoside-induced hair cell damage (Alassaf and Halloran, 2021), suggesting a role for growth factor signaling in protecting hair cells from damage.

While the activation of ERK and Akt signaling was similar to what was seen in other forms of cadmium-induced cell death, the fact that inhibiting these pathways increased rather than protected against cell death was unexpected. This, however, is not unprecedented. While many studies have found blocking Akt or ERK activation protects cells against cadmium-induced cell death (Chen et al., 2008; Iryo et al., 2000; Jiang et al., 2015; J. Kim et al., 2005; Zhang et al., 2017) others have found that it has no effect (Låg et al., 2005; Liu and Templeton, 2007; Rigon et al., 2008; Wang et al., 2008), and still others that death is increased similar to what we observed (Chao and Yang, 2001; Chuang et al., 2000; Xiao et al., 2009). Some of this disparity may come from differences in treatment paradigms. For example, in astrocytes ERK inhibition protects against chronic but not acute cadmium treatment (Jiang et al., 2015). The type of cell death also appears important as in macrophages ERK inhibition can protect against necrosis but not apoptosis induced by cadmium (J. Kim et al., 2005).

Unlike what we found in calcium signaling, the role of the Akt and ERK signaling pathways in cadmium-induced hair cell death appears similar to what has previously been shown for other forms of hair cell damage. These pathways are activated in response to both noise and other ototoxic drugs (Chung et al., 2006; Herranen et al., 2018; Ingersoll et al., 2020; Lahne and Gale, 2008; Maeda et al., 2013), and inhibiting them has been shown to increase hair cell death (Battaglia et al., 2003; Brand et al., 2015; Chung et al., 2006; Haake et al., 2009; Hayashi et al., 2013; Kurioka et al., 2015). Particularly striking is the fact that ERK activation is seen not in the hair cells themselves but rather in the support cells. This has also been seen in the mammalian inner ear in response to aminoglycosides, cisplatin, and noise (Herranen et al., 2018; Ingersoll et al., 2020; Lahne and Gale, 2008; Maeda et al., 2013).

Cadmium is known to damage support cells (Ding et al., 2020), and thus this ERK activation could be coming from a direct effect of cadmium on the support cells. Alternatively, the damaged hair cells could be sending a signal to the support cells to induce ERK activation. It has been shown that the damage-induced release of ATP and gap junction signaling are important for support cell ERK activation following mechanical injury of hair cells (Lahne and Gale, 2008). Once ERK is activated in support cells it may mediate its protective effect by modulating support cell released HSP70. ERK activation is important for HSP70 expression and folding (Lim et al., 2019; Xu et al., 2018; Zhai et al., 2019). Support cells are known to release HSP70 in exosomes following heat shock, with that HSP70 subsequently protecting hair cells from aminoglycoside-induced hair cell death (Breglio et al., 2020; May et al., 2013). The mechanism by which Akt activation is working in hair cell death is likely more straightforward as it is activated in the hair cells themselves. The activation of Akt signaling is well known to promote cell survival and inhibit cell death and multiple apoptotic regulators are Akt substrates (reviewed in Downward, 2004; Franke et al., 2003), providing a potential mechanism by which Akt activation could protect hair cells from cadmium-induced cell death.

The highly conserved activation and potential protective nature of Akt and ERK pathways suggest enhancing their activation could be a potential target for protecting hair cells from damage, however, as with other forms of cadmium-induced cell death, there are other examples where the activation of ERK appears to promote hair cell death (Lahne and Gale, 2008; So et al., 2007). The reasons behind these dual roles of ERK would need to be further investigated before ERK activation could be used as a potential therapeutic target.

## ACKNOWLEDGEMENTS

This work was supported by a Lafayette College Academic Research Committee faculty research grant to TMS and the Lafayette College Neuroscience program. We thank Amy Badillo for assistance with zebrafish care.

## AUTHOR CONTRIBUTIONS

**Jennifer Galdieri**: investigation, writing – original draft. **Chloe Adams**: investigation, writing – review & editing. **María Padilla**: investigation, writing – review & editing. **Tamara Stawicki**: conceptualization, methodology, formal analysis, investigation, writing – original draft and revision, and review and editing.

